# MSI.EAGLE: An Open-Source GUI for Streamlined Mass Spectrometry Imaging Analysis

**DOI:** 10.1101/2025.09.03.673811

**Authors:** Daniel J. Boehmler, Ashley Woolfork, Ai Wen Tan, Arjun Sengupta, Pinky Kain, Om B. Patel, Farheen Akhtar, George McClung, Daniel Ackerman, Terence P. Gade, Garret A. Fitzgerald, Aalim M. Weljie

## Abstract

Mass Spectrometry Imaging (MSI) is an emergent tool for analyzing spatial molecular distributions, yet data complexity often hinders effective analysis. MSI.EAGLE, an open-source R-Shiny application, makes analysis accessible for non-specialists by integrating advanced tools in a user-friendly interface. The workflow leverages tools from the Cardinal MSI package, with enhanced phenotyping, segmentation, statistical analysis and visualization. The application addresses a gap in spatial biology research by empowering a broader scientific community.

## MAIN TEXT

Mass Spectrometry Imaging (MSI) has emerged as a transformative technology for visualizing the spatial distribution of biomolecules, such as metabolites and biomarkers, directly from tissue samples.^1–3^ Current methodologies, including popular software packages like the Cardinal R package or commercial offerings, often present limitations in terms of user accessibility (i.e., the degree of user expertise required, cost, or both). While the data-intensive nature of MSI means that analysis is inherently complex, there is a lack of low-cost and simple-to-use user tools for analysis, which are crucial for the continued adoption of spatial lipid annotation and metabolic mapping.^4–6^

To bridge this gap, we introduce MSI.EAGLE, a sophisticated yet user-friendly R Shiny application designed for metabolic segmentation, phenotyping, and statistical analysis of mass spectrometry images. It has been built to leverage the strengths of MSI for both high-resolution tissue imaging and high-throughput sample extract screening. In our lab, DESI-MS, known for its minimal sample preparation requirements and ambient analysis capabilities^1,2,5^, forms the backbone of our approach; however, the tool is equally useful for MALDI-MS analysis (**Fig. S1)**. MSI.EAGLE integrates advanced data processing and statistical analysis provided by the Cardinal package^7^ via an intuitive graphical user interface (GUI) based workflow for researchers across various omics fields. This work outlines the development and application of MSI.EAGLE, demonstrating its potential to simplify and enhance MSI data analysis, facilitating access to the field of molecular imaging for non-specialists.

The MSI.EAGLE app is engineered to provide an intuitive, user-friendly workflow, via a a suite of advanced features suitable for complete analysis of MSI data. The app uses an R Shiny modular approach which allows for integration of additional modules and customization. Key features include project initialization with parallelization, robust data import options, efficient segmentation methods, masking of isolated pixel regions, image visualization and analysis tools, annotation, and statistical analysis capabilities (**Fig. 1**).

**Figure 1:**
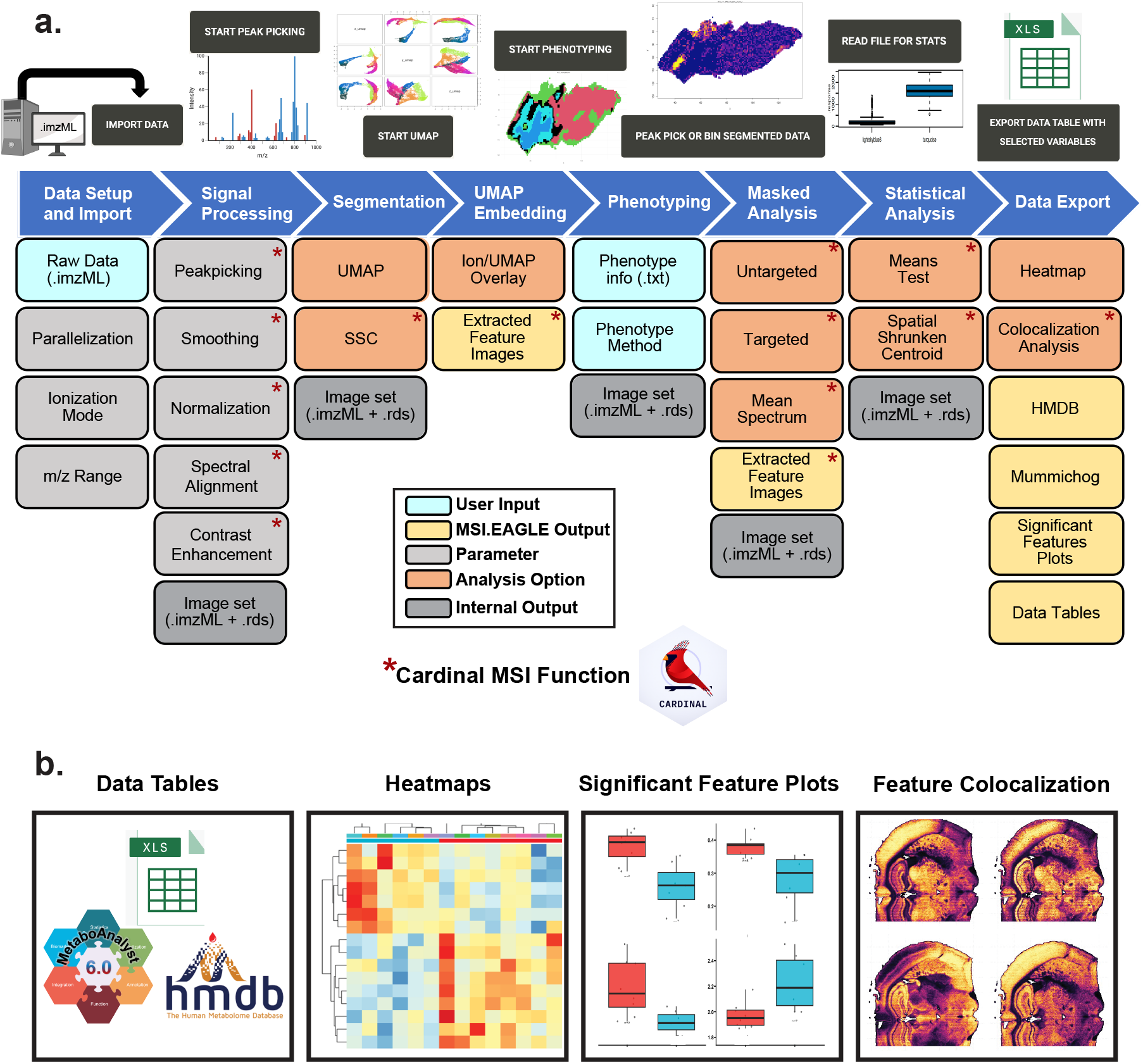
Workflow and capabilities of the MSI.EAGLE GUI. **a**. Analytical pipeline for processing and interpretation of mass spectrometry imaging (MSI) data. The workflow progresses through eight sequential modules: data import of raw imzML files, signal processing including peak picking and normalization, spatial segmentation, dimensionality reduction via UMAP embedding, phenotype definition, masking of regions of interest, statistical analysis, and data export. Color-coding indicates functional classifications: user inputs (blue), MSI.EAGLE outputs (yellow), adjustable parameters (gray), analysis options (orange), and internal outputs (gray). Interactive entry points specifying each module in the top section (black buttons) allow users to initiate key analytical processes. Red asterisks denote functions that leverage the Cardinal MSI package. **b**. Standardized outputs generated by MSI.EAGLE for downstream analysis and interpretation, including structured data tables for secondary analysis, interactive heatmaps for metabolite abundance visualization, significant feature plots highlighting statistical differences between regions, and feature colocalization maps for spatial correlation analysis. The integrated platform facilitates cohesive progression from raw MSI data to biologically meaningful features through a unified interface, enabling systematic tissue characterization and biomarker discovery.

The application consists of nine main functional modules: (1) **Data Setup** for file import and initial peak picking, (2) **File Restore and Overview Analysis** for data management and visualization, (3) **UMAP Segmentation** for dimensionality reduction-based clustering, (4) **UMAP Embedding** for interactive visualization of reduced-dimensional data, (5) **Spatial Shrunken Centroids (SSC) Segmentation** for spatially-aware clustering, (6) **Phenotyping** for associating metadata with samples, (7) **Masked analysis** for targeted feature extraction, (8) **Statistical Analysis** including means testing and spatial methods, and (9) **Heatmap and Outlier Detection** for data quality control and visualization.

To demonstrate the utility of MSI.EAGLE in analyzing locoregional changes in tissue metabolism, we analyzed murine brain tissue (**Fig 2a, 2b, 2c)**.^8^ Using the GUI for all steps, raw data were imported as .imzml format files, and initial peak-picking was performed within a specific mass range (*m/z* 200-300) to differentiate tissue from background. UMAP clustering^9^ was carried out on this data subset, allowing for the visualization of distinct clusters representing brain and background. This allowed for the selective removal of background pixels in the GUI. This step was crucial in isolating the brain tissue, allowing for a more targeted and meaningful analysis while dramatically reducing the computation burden of further analysis. Spatial Shrunken Centroids (SSC) segmentation can also be applied for this same purpose.^10^

**Figure 2:**
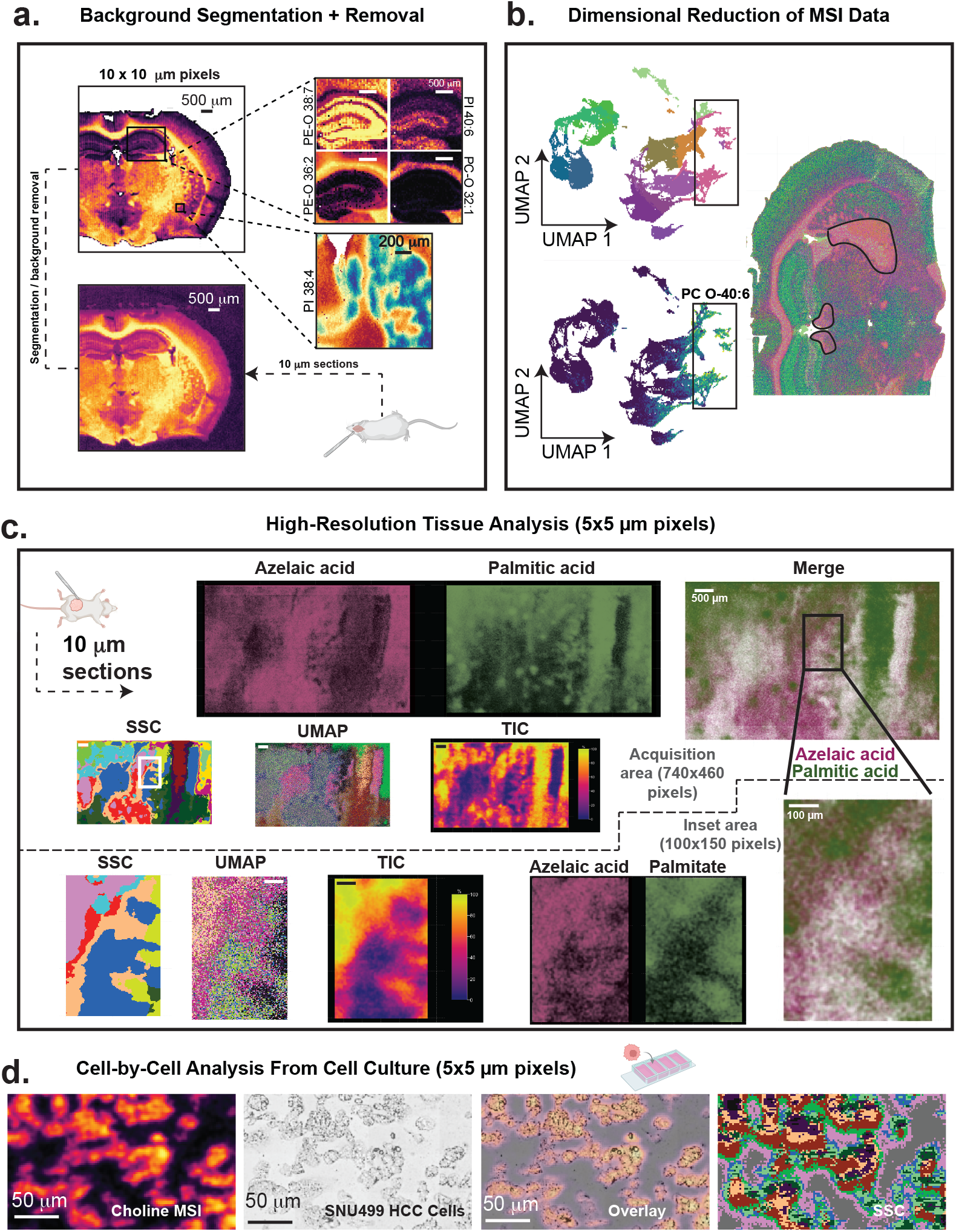
Application of MSI.EAGLE to mouse brain tissue and single-cell MSI analysis. MSI.EAGLE enables multi-scale mass spectrometry imaging analysis and data interpretation. **a**. Processing workflow showing segmentation and background removal of a mouse brain section imaged at 10 µm spatial resolution obtained via a click-through user interface. Insets highlight specific phospholipid distributions (PE-O 38:7, PE-O 36:2, PI 40:6, PC-O 32:1, PI 38:4) with differential abundance across hippocampal regions. **b**. UMAP dimensionality reduction of MSI data reveals distinct metabolic signatures corresponding to anatomical brain regions. UMAP embeddings on the left show clustering of similar biochemical profiles and one projected feature (PC-O 40:6) while the right displays the spatial projection of these clusters onto the original tissue section. **c**. Regional and highly localized DESI-MSI data imaged at 5 µm spatial resolution. Panels show HCC tumor tissue with SSC and UMAP clustering alongside corresponding MSI data in negative mode, revealing metabolic heterogeneity within tissues, as well as cell or cell neighborhood regionality. **d**. Single-cell DESI-MSI data imaged at 5 µm spatial resolution in positive mode overlaid with brightfield microscopy images at 40X. This analytical framework facilitates comprehensive interpretation of mass spectrometry imaging data across multiple biological scales, from whole-organ analysis to single-cell resolution.

Targeted analysis was performed to extract mass-to-charge ratios of known lipid species found by MSI in brain tissue through the **Masked Analysis** tab in MSI.EAGLE. Coordinates from the previously segmented datasets were mapped onto the raw data as the mask, which was re-imported with a mass range encompassing the entire targeted list (*m/z* 50-1000). This step revealed differences in metabolite abundances across specific brain substructures – demonstrated most strikingly in the hippocampus (**Fig. 2a)**. The final phase involved utilizing MSI.EAGLE’s GUI for phenotyping and statistical analysis. The UMAP clustering tool was revisited to annotate distinct brain regions, categorizing them based on lipid populations, which can then be mapped back onto UMAP plots (**Fig 2b)**. Post-analysis, information can then be exported in several formats such as tables compatible with HMDB and MetaboAnalyst queries, heatmaps, and ion images for presentation or publication.

As a further example of understanding metabolic heterogeneity at the cellular level, we demonstrate high spatial resolution MSI data processed with MSI.EAGLE (**Fig. 2c, 2d)**. In HCC tumor tissue^11^, MSI data were collected with 5 µm spatial resolution and then clustered via SSC and UMAP. SSC data reveal regionalized differences, characterized by complementary distribution of fatty acids such as palmitate and azelaic acid. Palmitate is higher in regions of active anabolic tumor activity, while azelaic acid is an end product of peroxidation from polyunsaturated acids such as linoleic/oleic acids following exposure to highly oxidative environments. Expansion of this data demonstrates that this heterogeneity is present in highly localized microenvironments, potentially in single cell or cell neighborhoods (**Fig. 2c)**.^12^ Additionally, in SNU449 HCC cells cultured in a monolayer on a microscope slide^13^, single cells were resolved in MSI data, then mapped to a brightfield microscopy image of the same cells (**Fig. 2d)** demonstrating intracellular heterogeneity via SSC analysis.

MSI can also be used for high-throughput chemical analysis.^14,15^ We demonstrate how MSI.EAGLE enhances accessible high-throughput processing capabilities through another application: characterizing the lipidomic profile of spotted human plasma extracts. (**Fig. S2)**. This use case addresses a significant challenge for researchers with limited computational expertise using open-source analysis tools for mass spectrometry imaging data, namely, the accessibility of efficient batch processing for multiple samples.^16^ While commercial platforms offer high-throughput solutions, our approach aims at low-cost, open-source, and readily accessible tools to improve reproducibility and throughput. In the GUI, 16 raw data files containing 578 samples were imported simultaneously at the full acquired mass range (*m/z* 50-1700), then batch segmented via UMAP clustering to isolate spots from background ionization. Spots were assigned specific phenotypes and replicates using an imported spreadsheet specific to the data set. Masking was performed by extracting mass-to-charge ratios of known lipid species on all files. Finally, means testing through the **Statistical Analysis** tab was performed to determine lipidomic changes between patient plasma samples.^17^ With our batch processing approach, this entire process can be accomplished on a computer with standard RAM capabilities (a Macintosh with 32 GB of RAM and M2 chip) in 15 minutes with no prior R experience (**Table S1)**. When files are processed individually, the time required for analysis increases exponentially, while processing outside the GUI requires substantial knowledge of both the Cardinal package and R. With our modular batch approach, laboratories processing large volumes of MSI data can increase data throughput many times.

In this communication, we introduce MSI.EAGLE, a comprehensive R Shiny application designed to meet the needs of application specialists in the analysis of Mass Spectrometry Imaging (MSI) data. MSI.EAGLE represents a leap forward in MSI data processing and analysis. Its development aligns with the growing need for user-friendly, robust, customizable, and efficient tools in the rapidly evolving field of mass spectrometry imaging. As MSI data continue to grow in complexity and volume, such tools will be indispensable for researchers to unlock the full potential of MSI technology. Installation instructions and source code are available for access here: https://github.com/Weljie-Lab-UPenn/MSI.EAGLE

## ONLINE METHODS

### Sample Preparation

We employed a biphasic extraction protocol for isolating polar metabolites and lipids from aqueous serum samples.^18,19^ For high-throughput screening, 1 µL of each sample extract was spotted onto PTFE-coated slides for DESI-MS analysis. For tissue imaging, flash-frozen samples were embedded in 12.5% fish gelatin, sectioned at 10 µm thickness and mounted on superfrost glass microscope slides, and stored at −80°C until analysis.^5,20,21^ For cell culture samples, cells were plated in 4-well cell culture slides (Part No: CCS-4, MatTek) at 500K cells per well and incubated in DMEM +10% FBS with Glutamax for 24 hours to adhere to the slide surface. Media was removed and wells washed with PBS before drying and MSI acquisition.

### DESI-MS System Setup and Data Acquisition

DESI-QTOF mass spectrometry imaging data were acquired using a Xevo G2-XS QTOF mass spectrometer equipped with a DESI-XS source (Waters Corporation, Milford, MA, USA). The mass spectrometer was operated in both negative and positive ion modes. In negative ion mode, the spray solvent consisted of 98% methanol with 50 pg/mL leucine enkephalin as an internal standard. For positive ion mode, the spray solvent was composed of 98% methanol with 50 pg/mL leucine enkephalin and 0.01% formic acid. The flow rate of the spray solvent was set to 350 nL/min for < 5 × 5 μm pixel data, 1 μL/min for > 5 × 5 μm pixel data in both modes.

The DESI source parameters were optimized for each ion mode. The spray voltage was set to −0.5 kV in negative mode and +0.9 kV in positive mode. The nebulizing N_2_ gas flow was maintained at 600 L/H and the heated transfer line temperature was set to 50°C for mouse kidney and SNU449 cell acquisition, and 400°C for spotted samples, mouse brain, and drosophila brain tissue. The sampling area was defined using High Definition Imaging (HDI) software, and the pixel size varied from 2 μm in cells and tissues to 200 μm for spotted plates. The mass spectrometer was operated in sensitivity mode with a mass range of m/z 50-1200.

### Data Extraction and Analysis

RAW data were converted to .mzml and .imzml formats using MSConvert GUI (ProteoWizard) and imzML Converter 1.3, respectively.^6^ The processed imzML files were then imported into R for further visualization and analysis using MSI.EAGLE.

### Data Import and Setup

The data import and setup tab provides users with functionality to import, visualize, and preprocess MSI datasets. MSI.EAGLE users can specify the directory containing raw imzML files and a working directory for result storage within the data import section. Files are selected through an interactive data table in the GUI. This step supports parallel processing, with users able to configure the parallelization mode and number of cores.

For peak picking, two predefined parameter sets are provided. “qTof1” is suitable for quadrupole time-of-flight instruments, while “HiRes” is designed for high-resolution instruments. Users have the flexibility to adjust these parameters based on their specific acquisition instrument and analysis type (**Table S2**).

The data visualization module generates quick visualizations of the imported data, allowing users to verify acquisition and data conversion quality before proceeding to more in-masked analysis. A mass spectrum plot is created using a user-defined percentage of randomly sampled pixels, typically set to a low value. Users can interactively adjust the m/z and intensity ranges for this plot, helping them decide on relevant mass ranges for further analysis steps.

For testing purposes, the demo data section of this tab includes options to load pre-processed datasets from the CardinalWorkflows package. These datasets include samples such as human renal cell carcinoma and pig fetus cross-sections.

The data setup module is built on reactive programming principles in Shiny, ensuring efficient updates to outputs based on user inputs. This approach optimizes the application’s responsiveness by performing computationally intensive operations, like data import and peak picking of large datasets, only when necessary.

### File Restore and Overview Analysis

The sidebar panel allows users to select from four main operations: 1) opening a previously peak-picked file, 2) peak picking raw files, 3) adding two imagesets with the same coordinates, or 4) adding two imagesets with the same peak list. These options enable users to choose the appropriate workflow for the stage and structure of their data.

For de novo peak picking, users can specify parameters such as the percentage of pixels to sample, signal-to-noise ratio (SNR), minimum peak frequency, and peak picking method (simple, adaptive, or mad). The app utilizes Cardinal’s peakPick() function with these user-defined parameters to identify peaks in the MSI data.

The main panel displays a dynamic table of the processed runs, allowing users to select specific runs for further analysis. An interactive MSI plot is generated, which provides a visual representation of the peak-picked data. The module also includes functionality for saving the processed data as a .imzML directory and RDS file. This ensures that users can easily export their processed data for further analysis or future use, avoiding arduous re-processing steps.

### Segmentation – Uniform Manifold Approximation and Projection (UMAP)

MSI.EAGLE implements unsupervised segmentation of mass spectrometry imaging data using Uniform Manifold Approximation and Projection (UMAP) dimensionality reduction.^9^ Users can choose between background removal based on pixel data or anatomical segmentation to generate phenotype data. The UMAP algorithm is applied to the selected dataset using the uwot R package.^22^ Key parameters including minimum distance, number of neighbors, and number of trees can be adjusted through the GUI.

UMAP analysis is performed on the pixel intensity data, and the resulting low-dimensional UMAP embeddings are then clustered using one of ten algorithms. K-means is the default method, but options include: hierarchical, DBSCAN, HDBSCAN, Spectral Clustering, K-medoids, Fuzzy C-means, Model-based Clustering (Mclust), Self-Organizing Map (SOM), and Spherical K-means. Users can interactively visualize the UMAP results and clustering, with options to display reduced color representations or cluster assignments. The GUI allows selection of specific clusters or colors to refine the segmentation.

Processed segmentation results can be stored and applied to the full dataset. For anatomical segmentation, users can annotate specific regions by selecting clusters and assigning labels. The app provides additional functionality to remove isolated pixels to clean up the segmentation. Processed and annotated data can be saved for downstream analysis as both an .imzML directory and .rds file in the working directory.

The segmentation tab leverages the Cardinal package for handling and visualizing mass spectrometry imaging data structures. Interactive plots are generated using ggplot2 and rendered as PNG images for presentation and/or publication by the user. This segmentation approach enables flexible, data-driven partitioning of imaging mass spectrometry data without relying on predefined anatomical boundaries. The interactive interface also allows iterative refinement of segmentation results, introducing tolerance for sub-optimal parameter selection by the user.

### Segmentation – Spatial Shrunken Centroids (SSC)

MSI.EAGLE also utilizes the spatialShrunkenCentroids function from the Cardinal package to perform unsupervised segmentation of MSI data. This approach, based on the shrunken centroid method, incorporates spatial information to identify regions with similar molecular profiles.

The GUI allows users to select preprocessed datasets for analysis and set key parameters for the spatialShrunkenCentroids algorithm, including the spatial neighborhood radius (r), number of clusters (k), and shrinkage factor (s). Multiple parameter combinations can be evaluated in parallel. The segmentation results are cached to avoid redundant computations. Again, there is an optional step to remove isolated pixels is available through the fix_pix function to clean the image data.

Segmentation results are visualized using Cardinal’s image and plot functions. The GUI provides options to select specific segmentation models and color schemes for visualization. Users can interactively explore different cluster assignments and evaluate corresponding weighted spectra and statistics. The module allows storing and exporting the processed data, including segmentation results, for further analysis or reporting. All processing steps maintain the spatial and spectral integrity of the imaging data structure as implemented in Cardinal.

This segmentation approach enables users to objectively differentiate background ionization from sample data and identify regions of interest in mass spectrometry imaging experiments, facilitating the discovery of spatially-resolved molecular patterns in complex biological samples.

### UMAP Embedding

MSI.EAGLE’s UMAP Embedding module enables interactive visualization and exploration of mass spectrometry imaging data in reduced dimensional space. The module utilizes a React-based interface for real-time interaction with UMAP projections. Users can select m/z ions of interest and color variables from pixel metadata through a dropdown menu interface. The embedding visualization supports both intensity-based coloring of selected m/z values and categorical coloring based on metadata variables.

For intensity-based visualization, the module implements a color scaling system with an adjustable midpoint value and optional log transformation. Points in the UMAP space can be selectively masked based on metadata values, with unselected points appearing in a neutral gray. Point size and other visual parameters can be adjusted through interactive controls. A complementary spatial plot shows the selected m/z value or metadata variable mapped back onto the original image coordinates. All visualization options can be exported as high-resolution images for presentation or publication.

### Phenotyping

The phenotyping tab of MSI.EAGLE enables users to associate phenotypic data with MSI datasets. Users can upload a tab-delimited text file containing phenotype information via a file input interface. The uploaded file is read using the read_samples function, which automatically detects the file format.

Users can choose between two types of MSI images for phenotyping: 1) images restored from a .imzml or .rds file or 2) stored data from the application memory.

The application offers four phenotyping methods: 1) Spectral density, 2) Periodicity, 3) Breaks between samples, and 4) Manual (x & y limits specified in file). For the “breaks” method, users can specify a threshold value. The pixDatFill_mult function is used to perform the phenotyping for the first three methods, while pixDatFill_manual is used for the manual method.

After phenotyping, the results are displayed in an interactive table using the DT package. Users can select which columns from the phenotype data to include in the pData object. The application also provides an option to create interaction terms between two selected phenotype variables. The phenotyped data is visualized using Cardinal’s image function, allowing users to select which phenotype variable to display. The resulting image is rendered as a PNG file and displayed in the user interface. Users can save the phenotyped data, which includes the MSI image data with updated pData and fData objects.

This phenotyping module provides a user-friendly interface for associating phenotypic data with MSI datasets, enabling researchers to explore relationships between molecular profiles and sample characteristics within the application.

### Masked Analysis

The Masked Analysis tab allows users to perform more comprehensive peak picking and extraction of lower abundance features. Users first select a previously segmented dataset file as a template for coordinate extraction. Coordinate extraction is performed by iterating over the run names in the segmented file and extracting corresponding coordinates using Cardinal’s coord function. The module then offers three analysis modes: untargeted peak picking, targeted peak binning, and mean spectrum-based peak picking.

For untargeted analysis, peak picking is performed using a custom HTS_reproc function, which incorporates multiple Cardinal functions. Targeted analysis utilizes Cardinal’s peakBin function, allowing users to input a list of annotated exact masses for binning. The mean spectrum approach combines Cardinal’s summarizeFeatures, normalize, peakPick, peakAlign, and peakFilter functions to calculate an average spectrum, perform peak picking, and use the resulting peak list to bin the raw data. Parallel processing capabilities are incorporated to handle large datasets, utilizing the bplapply function for certain operations.

After processing, the resulting peak-picked or binned data is visualized using Cardinal’s image function. The processed data can be saved for further analysis. This functionality enables researchers to extract and analyze depth-specific molecular information from complex samples in an interactive and user-friendly manner.

### Statistics

The Statistics tab enables users to perform various statistical tests on the processed MSI data. The GUI allows selection of the analysis type, data grouping, and visualization options. Statistical methods implemented include means tests and spatial shrunken centroids (SSC).

For means tests, the app utilizes Cardinal’s meansTest function to compare ion intensities between user-defined groups. Results are displayed as an interactive table with adjustable false discovery rate (FDR) thresholds. The spatial method (SSC) incorporates pixel coordinates to identify spatially-localized differences between groups.

MSI.EAGLE generates customizable visualizations of statistical results, including boxplots, means plots, and ion images colored by group membership or statistical significance. Users can interactively explore results, adjust plot parameters, and export tables and figures.

The modular design and open-source nature of MSI.EAGLE allows easy addition of new statistical tests and visualization options. All analyses are performed server-side using R, with results passed to the user interface for display and interaction. This architecture enables efficient analysis of large imaging datasets within memory constraints.

### Heatmap and Outlier Detection

MSI.EAGLE provides interactive heatmap visualization and outlier detection capabilities through a dedicated tab. Users can select variables for heatmap generation from phenotypic data associated with the MSI dataset. The heatmap is generated using the pheatmap R package, with options for row/column scaling, clustering methods, distance metrics, and color schemes.

Significance filtering can be applied to show only features meeting a user-defined threshold based on statistical results. Row and column annotations can be added to provide additional context. The resulting heatmap can be customized through various parameters including clustering cutoffs, label display, and color palettes.

Outlier detection is implemented using the stray R package, which applies a high-dimensional outlier detection algorithm. Users can adjust key parameters such as the number of nearest neighbors (k), significance level (alpha), and proportion of potential outliers. MSI.EAGLE identifies outlier samples and provides options to regenerate the heatmap with outliers removed or save a filtered dataset excluding detected outliers.

All visualizations can be exported as high-resolution PDF files. Associated data matrices (raw and scaled) are saved as tab-separated text files to facilitate further analysis. This functionality enables iterative exploration and quality control of high-dimensional mass spectrometry imaging data within an interactive framework.

### Colocalization Analysis

In the Colocalization Analysis tab of MSI.EAGLE, options include selecting correlation variables and m/z values, numeric inputs for specifying the number of colocalized features and top correlations to plot, and options for choosing the sorting method.

Upon triggering the analysis, the colocalization analysis is performed using the colocalized function from Cardinal. This analysis considers the user-selected m/z value, number of features, and sorting method. Visualization of the colocalization results is achieved using Cardinal’s image function. The resulting plot displays the top correlated features as specified by the user, with options for contrast enhancement, normalization, and color scale customization. This implementation allows for interactive exploration of colocalization patterns in mass spectrometry imaging data, providing users with flexible options for analysis and visualization.

### Co-Registration of H&E microscopy images with MSI data

Co-registration of mass spectrometry images with H&E microscope images of the same tissue section was performed outside of the MSI.EAGLE application. H&E images were acquired using a uSCOPE MXII microscope at 40X, then imported to QuPath for further processing. Using QuPath’s built-in cell detection algorithms, cells were automatically identified based on nuclear staining intensity and cell size parameters. The detected cells were manually verified and adjusted where needed. Cell region boundaries were exported as polygons from QuPath stored as GeoJSON files.

Co-registration of these polygons with MSI data was performed in an interactive interface. Both the GeoJSON file containing the QuPath annotations and the processed MSI data (.imzML or .rds format) were loaded. Registration was performed through an affine transformation combining translation, rotation, and scaling. The transformation parameters were specified manually through the interface, and the overlay transparency was adjusted to visually verify alignment quality.

This process maps individual cells onto the MSI data coordinates. The co-registered data was then exported as both .rds and .imzML files compatible with MSI.EAGLE, with the cell annotations preserved in the pixel metadata. Additionally, two tables were exported: a region measurements table containing morphological features extracted from QuPath for each cell (including area, perimeter, and circularity), and a transformation parameters table recording the specific scaling, rotation, and translation values used for registration to ensure reproducibility.

## Contributions

**D.B**., **A.W**., and **A.W.T**. contributed equally to this work. **D.B**. and **A.M.W**. developed the MSI.EAGLE software architecture, implemented the R-Shiny interface, and performed software testing. **D.B**. acquired mouse brain data, tumor data, and cell culture data. **D.B**. wrote the manuscript with input from all authors. **A.W**. contributed to software development, and prepared mouse brain tissue sections.

**A.W.T**. contributed to software development. **A.S**. collected spotted plasma sample data. **P.K**. assisted in manuscript preparation. **O.B.P**. contributed to software development. **F.A**. assisted in manuscript preparation. **G.M**. and **D.A**. implanted xenograft tumors in mice. **T.P.G**. supervised the tumor sample collection and analysis. **G.A.F**. assisted in manuscript preparation. **A.M.W**. conceived and supervised the project, providing funding. All authors reviewed and approved the final manuscript.

## Acknowledgments

This work was supported by grants from the National Institutes of Health: 1R01NR018836 from the National Institute of Nursing Research (NINR), 1R01HL142981 from the National Heart, Lung, and Blood Institute (NHLBI), 5R01DK120757 from the National Institute of Diabetes and Digestive and Kidney Diseases (NIDDK), and P01CA165997 from the National Cancer Institutes (NCI).

## SUPPLEMENTARY INFORMATION

**Figure S1:**
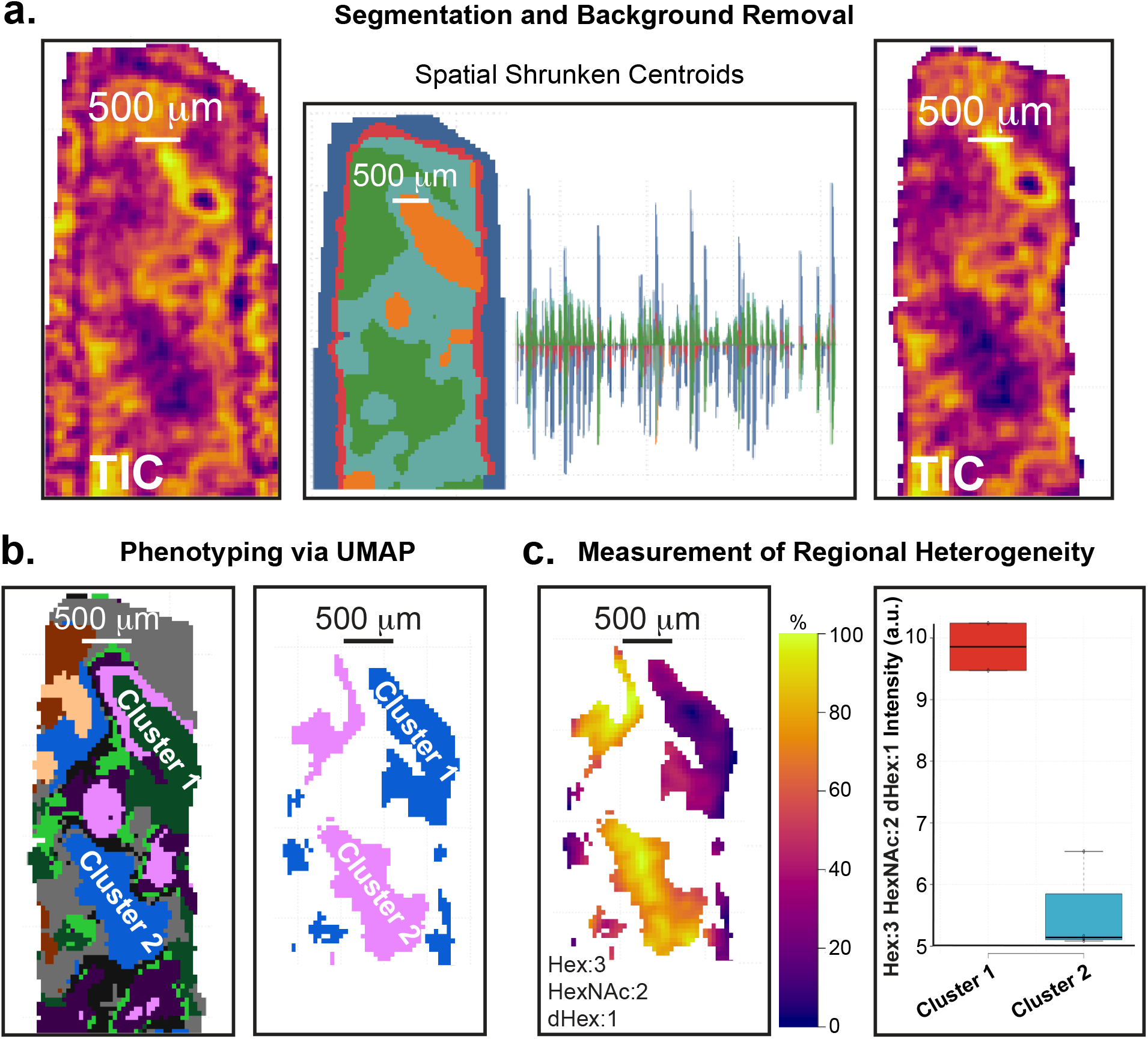
MSI.EAGLE processing of MALDI MSI data. **a**. Segmentation and background removal process using a publicly available MALDI-FTICR dataset of FFPE human kidney embedded in CHCA MALDI matrix with 50 × 50 µm pixels (https://metaspace2020.org/dataset/2024-11-09_00h04m19s). Raw MALDI-MSI images (top) were peak-picked at 10 ppm tolerance, then underwent spatial shrunken centroids segmentation, producing region-specific molecular signatures with corresponding spatial domains (bottom). **b**. Unsupervised phenotyping via UMAP dimensionality reduction reveals distinct molecular clusters. Left panel shows all identified clusters, while right panel demonstrates selective visualization of key clusters (Cluster 1 and Cluster 2) for subsequent metabolic analysis. **c**. Quantitative assessment of regional metabolic heterogeneity. Left panel displays spatial distribution of a specific N-glycan (Hex:3 HexNAc:2 dHex:1) across the tissue section. Right panel shows significant differential intensity of this glycan between Cluster 1 and Cluster 2, revealing metabolic subpopulations within the tissue microenvironment. The untargeted differential analysis approach enables discovery of region-specific metabolic features without prior molecular targeting.

**Figure S2:**
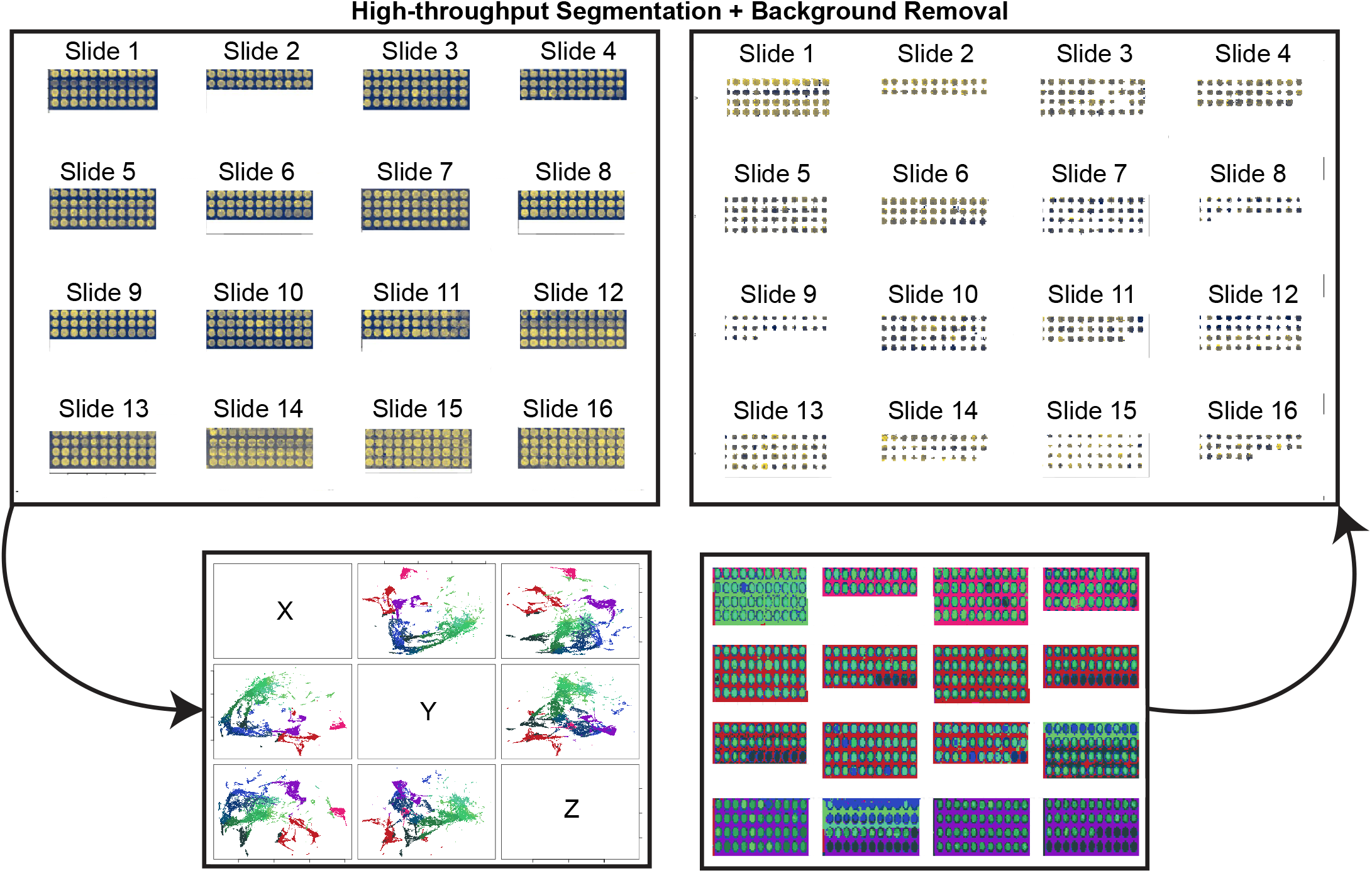
Application of MSI.EAGLE to high-throughput plating experiments. A demonstration of the high-throughput capabilities of MSI.EAGLE through batch processing of spotted human plasma extracts. This demonstrates parallel peak-picking, segmentation, background removal, and phenotyping based on pixel location for multiple sample slides simultaneously. This high-throughput analytical approach enabled the comprehensive lipidomic analysis of plasma samples from 149 subjects (67 COVID-19 patients, 66 non-COVID-19 sepsis patients, and 16 healthy controls), facilitating the identification of COVID-19-specific lipid signatures. The batch processing capabilities were essential for discovering the three high abundance lipids (ChoE 18:3, LPC-O-16:0, and PC-O-30:0) that showed specific alterations in COVID-19 patients compared to other sepsis causes and healthy controls, and which correlated with disease severity.

**Table S1:** Data and Analysis Characteristics.

| Data Characteristic | HCC Patient-Derived Xenograft (Fig. 1a) | Mouse Brain (Fig. 2a, 2b) | HCC Patient-Derived Xenograft ROI (Fig. 2c) | SNU449 HCC Cells (Fig. 2d) | Human Plasma Spots (Fig. S2) | MALDI Human Lung (Fig. S1) |
| --- | --- | --- | --- | --- | --- | --- |
| Processing time (minutes from data import to images shown in figures)** | 3.5 | 20 | 60 | 90 | 15 | 10 |
| Number of features detected after untargeted peak-picking (on-sample, 30 ppm window, 20 S/N) | 5,988 | 1,424 | 5,791 | 4,252 | 622 (targeted) | 94* |
| Pixel size (μM) | 100 x 100 | 10 x 10 | 2 x 2 | 2 x 2 | 200 x 200 | 50 x 50 |
| Estimated mass resolution (ppm) | 6.5 | 9.5 | 17 | 11.5 | 5 to 7 | 4 |
| Number of pixels | 82152 | 817500 | 250000 | 1000000 | 320000 | 5459 |
| Raw .imzML file size (Gb) | 0.3 | 2.8 | 0.9 | 3.4 | 0.2 | 0.01 |
| Raw .ibd file size (Gb) | 6.8 | 40.5 | 4.8 | 30.9 | 7.47 (16 slides) | 0.1 |

\*annotated features from targeted peak-picking list
\*\*All processing done on M1 Macbook Pro with 32 Gb RAM

**Table S2:** Tabulated Features of MSI.EAGLE By Sample Analysis.

| MSI.EAGLE Tab | Parameter | HCC Patient-Derived Xenograft (Fig. 1a) | Mouse Brain (Fig. 2a, 2b) | HCC Patient-Derived Xenograft ROI (Fig. 2c) | SNU449 HCC Cells (Fig. 2d) | Human Plasma Spots (Fig. S2) | MALDI Human Lung (Fig. S1) |
| --- | --- | --- | --- | --- | --- | --- | --- |
| DATA SETUP | Parallelization mode | MulticorParam | MulticorParam | MulticorParam | MulticorParam | MulticorParam | MulticorParam |
| DATA SETUP | Parallelization mode | MulticorParam | MulticorParam | MulticorParam | MulticorParam | MulticorParam | MulticorParam |
| DATA SETUP | Number of cores to use | 8 | 8 | 8 | 8 | 8 | 8 |
| DATA SETUP | Peak picking params | qTof1 | qTof1 | qTof1 | 50 | qTof1 | Hi Res |
| DATA SETUP | Resolution for peakpicking (ppm) | 15 | 15 | 15 | qTof1 | 15 | 10 |
| DATA SETUP | Tolerance for spectral alignment (ppm) | 50 | 50 | 50 | 30 | 50 | 50 |
| DATA SETUP | m/z min for import | 200 (segmentation) and 50 (masked analysis) | 750 (segmentation) and 50 (masked analysis) | 200 (segmentation) and 50 (masked analysis) | 30 | 50 | 1000 |
| DATA SETUP | m/z max for import | 300 (segmentation) and 1700 (masked analysis) | 800 (segmentation) and 1200 (masked analysis) | 300 (segmentation) and 1200 (masked analysis) | 750 | 1700 | 3000 |
| DATA SETUP | Contrast enhancement | histogram | histogram | histogram | (segmentation) and 50 (masked analysis) | histogram | histogram |
| DATA SETUP | % of pixels to randomly sample for plot | 1 | 1 | 1 | 1 | 1 | 1 |
| DATA SETUP | Contrast enhancement | histogram | histogram | histogram | histogram | histogram | histogram |
| DATA SETUP | Colorscale | plasma | inferno | inferno/purple-green | inferno | cividis | inferno |
| DATA SETUP | Smoothing options | none | gaussian | gaussian | gaussian | none | none |
| FILE RESTORE AND OVERVIEW ANALYSIS - Peakpick rawfiles (imzML loaded in the Data Setup Tab) | % of pixels to randomly sample for peakpicking | 1 | 1 | 1 | 1 | 1 | 1 |
| FILE RESTORE AND OVERVIEW ANALYSIS - Peakpick rawfiles (imzML loaded in the Data Setup Tab) | S/N for overview peakpicking | 20 | 20 | 20 | 20 | 20 | 20 |
| FILE RESTORE AND OVERVIEW ANALYSIS - Peakpick rawfiles (imzML loaded in the Data Setup Tab) | Minimum peak frequency (0-1) | 0.01 | 0.01 | 0.01 | 0.01 | 0.01 | 0.01 |
| FILE RESTORE AND OVERVIEW ANALYSIS - Peakpick rawfiles (imzML loaded in the Data Setup Tab) | Peak picking method | diff | diff | mad | diff | mad | diff |
| SEGMENTATION - UMAP - Background Removal | Minimum distance for UMAP | 0.1 | 0.1 | 1 | N/A | 0.1 | 0.1 |
| SEGMENTATION - UMAP - Background Removal | # of nodes to search | 50 | 50 | 50 | N/A | 50 | 50 |
| SEGMENTATION - UMAP - Background Removal | UMAP nearest neighbors | 5 | 5 | 15 | N/A | 5 | 5 |
| SEGMENTATION - UMAP - Background Removal | Number of trees for UMAP | 50 | 50 | 50 | N/A | 50 | 50 |
| SEGMENTATION - UMAP - Background Removal | set_op_mix_ratio | 1 | 1 | 0.1 | N/A | 1 | 1 |
| SEGMENTATION - UMAP - Background Removal | UMAP PCA components (NULL for no PCA) | N/A | N/A | N/A | N/A | N/A | N/A |
| SEGMENTATION - UMAP - Background Removal | Method for HDBSCAN calculation | R DBSCAN package | R DBSCAN package | R DBSCAN package | N/A | R DBSCAN package | R DBSCAN package |
| SEGMENTATION - UMAP - Background Removal | Tolerance for spectral alignment (ppm) | 0.15 | 0.15 | 0.15 | N/A | 0.15 | 0.15 |
| SEGMENTATION - UMAP - Background Removal | DBSCAN eps | 50 | 50 | 50 | N/A | 50 | 50 |
| SEGMENTATION - UMAP - Background Removal | H/DBSCAN min points / cluster | 50 | 50 | 50 | N/A | 50 | 50 |
| SEGMENTATION - UMAP - Background Removal | cex, tile plot size adjustment | 2 | 2 | 2 | N/A | 2 | 2 |
| SEGMENTATION - UMAP - Background Removal | R color reduction quantile (0-1) | 0.9 | 0.9 | 0.9 | N/A | 0.9 | 0.9 |
| SEGMENTATION - UMAP - Anatomical Segmentation | Minimum distance for UMAP | N/A | N/A | N/A | N/A | N/A | N/A |
| SEGMENTATION - UMAP - Anatomical Segmentation | # of nodes to search | N/A | N/A | N/A | N/A | N/A | N/A |
| SEGMENTATION - UMAP - Anatomical Segmentation | UMAP nearest neighbors | N/A | N/A | N/A | N/A | N/A | N/A |
| SEGMENTATION - UMAP - Anatomical Segmentation | Number of trees for UMAP | N/A | N/A | N/A | N/A | N/A | N/A |
| SEGMENTATION - UMAP - Anatomical Segmentation | set_op_mix_ratio | N/A | N/A | N/A | N/A | N/A | N/A |
| SEGMENTATION - UMAP - Anatomical Segmentation | UMAP PCA components (NULL for no PCA) | N/A | N/A | N/A | N/A | N/A | N/A |
| SEGMENTATION - UMAP - Anatomical Segmentation | Method for HDBSCAN calculation | N/A | N/A | N/A | N/A | N/A | N/A |
| SEGMENTATION - UMAP - Anatomical Segmentation | Tolerance for spectral alignment (ppm) | N/A | N/A | N/A | N/A | N/A | N/A |
| SEGMENTATION - UMAP - Anatomical Segmentation | DBSCAN eps | N/A | N/A | N/A | N/A | N/A | N/A |
| SEGMENTATION - UMAP - Anatomical Segmentation | H/DBSCAN min points / cluster | N/A | N/A | N/A | N/A | N/A | N/A |
| SEGMENTATION - UMAP - Anatomical Segmentation | cex, tile plot size adjustment | N/A | N/A | N/A | N/A | N/A | N/A |
| SEGMENTATION - UMAP - Anatomical Segmentation | R color reduction quantile (0-1) | N/A | N/A | N/A | N/A | N/A | N/A |
| SEGMENTATION - UMAP - Fix stray pixels | Radius to search for neighbors | 3 | 3 | 3 | N/A | 3 | 3 |
| SEGMENTATION - UMAP - Fix stray pixels | min # of neighbors allowed | 2 | 2 | 2 | N/A | 2 | 2 |
| SEGMENTATION - SSC | Radius for SSC | 3 | 3 | 2 | 2 | 3 | 3 |
| SEGMENTATION - SSC | k (clusters) for SSC | 5 | 5 | 5 | 5 | 5 | 5 |
| SEGMENTATION - SSC | Shrinkage for Cardinal SSC | 2^(1:6) | 2^(1:6) | 2^(1:6) | 2^(1:6) | 2^(1:6) | 2^(1:6) |
| SEGMENTATION - SSC | Method for SSC | adaptive | adaptive | adaptive | adaptive | adaptive | adaptive |
| PHENOTYPING | MSI image to phenotype | Peakpicked file | Peakpicked file | Peakpicked file | Peakpicked file | Peakpicked file | Peakpicked file |
| PHENOTYPING | Phenotype method | N/A | N/A | N/A | N/A | Spectral density | N/A |
| MASKED ANALYSIS - Untargeted | S/N for comprehensive peak picking | N/A | N/A | N/A | N/A | N/A | N/A |
| MASKED ANALYSIS - Untargeted | Minimum peak frequency (0-1) | N/A | N/A | N/A | N/A | N/A | N/A |
| MASKED ANALYSIS - Untargeted | Tolerance for alignment or peak binning (ppm) | N/A | N/A | N/A | N/A | N/A | N/A |
| MASKED ANALYSIS - Untargeted | Peak picking method | N/A | N/A | N/A | N/A | N/A | N/A |
| MASKED ANALYSIS - Targeted | Tolerance for peak binning (ppm) | 15 | 15 | 15 | 30 | 15 | 15 |
| MASKED ANALYSIS - Mean Spectrum | S/N for comprehensive peak picking | N/A | N/A | N/A | N/A | N/A | N/A |
| MASKED ANALYSIS - Mean Spectrum | Peak picking method | N/A | N/A | N/A | N/A | N/A | N/A |
| MASKED ANALYSIS - Mean Spectrum | Minimum peak frequency (0-1) | N/A | N/A | N/A | N/A | N/A | N/A |
| MASKED ANALYSIS - Mean Spectrum | Tolerance for peak binning (ppm) | N/A | N/A | N/A | N/A | N/A | N/A |
| STATISTICS - Means test (Cardinal) | FDR threshold to include in results | N/A | N/A | N/A | N/A | N/A | 0.05 |
| STATISTICS - Means test (Cardinal) | Choose variable to test | N/A | N/A | N/A | N/A | N/A | Rcol_reduced |
| STATISTICS - Means test (Cardinal) | Choose members to include | N/A | N/A | N/A | N/A | N/A | UMAP Colors |
| STATISTICS - Means test (Cardinal) | Choose interaction for grouping | N/A | N/A | N/A | N/A | N/A | x |
| STATISTICS - Means test (Cardinal) | Choose variables for modeling | N/A | N/A | N/A | N/A | N/A | Rcol_reduced |
| STATISTICS - Spatial shrunken centroids (Cardinal) | FDR threshold to include in results | N/A | N/A | N/A | N/A | N/A | N/A |
| STATISTICS - Spatial shrunken centroids (Cardinal) | Choose variable to test | N/A | N/A | N/A | N/A | N/A | N/A |
| STATISTICS - Spatial shrunken centroids (Cardinal) | Choose members to include | N/A | N/A | N/A | N/A | N/A | N/A |
| STATISTICS - Spatial shrunken centroids (Cardinal) | SSC r value | N/A | N/A | N/A | N/A | N/A | N/A |
| STATISTICS - Spatial shrunken centroids (Cardinal) | SSC s value | N/A | N/A | N/A | N/A | N/A | N/A |
| STATISTICS - Spatial shrunken centroids (Cardinal) | SSC k value | N/A | N/A | N/A | N/A | N/A | N/A |
| STATISTICS - Spatial shrunken centroids (Cardinal) | Choose interaction for grouping | N/A | N/A | N/A | N/A | N/A | N/A |
| STATISTICS - Spatial shrunken centroids (Cardinal) | Choose variables for modeling | N/A | N/A | N/A | N/A | N/A | N/A |
| STATISTICS - Spatial DGMM (Cardinal) | FDR threshold to include in results | N/A | N/A | N/A | N/A | N/A | N/A |
| STATISTICS - Spatial DGMM (Cardinal) | Choose variable to test | N/A | N/A | N/A | N/A | N/A | N/A |
| STATISTICS - Spatial DGMM (Cardinal) | Choose members to include | N/A | N/A | N/A | N/A | N/A | N/A |
| STATISTICS - Spatial DGMM (Cardinal) | DGMM r value | N/A | N/A | N/A | N/A | N/A | N/A |
| STATISTICS - Spatial DGMM (Cardinal) | DGMM k value | N/A | N/A | N/A | N/A | N/A | N/A |
| STATISTICS - Spatial DGMM (Cardinal) | Choose interaction for grouping | N/A | N/A | N/A | N/A | N/A | N/A |
| STATISTICS - Spatial DGMM (Cardinal) | Choose variables for modeling | N/A | N/A | N/A | N/A | N/A | N/A |
| STATISTICS - ANOVA (Experimental) - One way (Y ~ A) | FDR threshold to include in results | N/A | N/A | N/A | N/A | N/A | N/A |
| STATISTICS - ANOVA (Experimental) - One way (Y ~ A) | Choose variable to test | N/A | N/A | N/A | N/A | N/A | N/A |
| STATISTICS - ANOVA (Experimental) - One way (Y ~ A) | Choose grouping variables | N/A | N/A | N/A | N/A | N/A | N/A |
| STATISTICS - ANOVA (Experimental) - One way (Y ~ A) | Choose variables for modeling | N/A | N/A | N/A | N/A | N/A | N/A |
| STATISTICS - ANOVA (Experimental) - One way (Y ~ A) | Y Variable for one-way ANOVA | N/A | N/A | N/A | N/A | N/A | N/A |
| STATISTICS - ANOVA (Experimental) - One way (Y ~ A) | Choose members to include | N/A | N/A | N/A | N/A | N/A | N/A |
| STATISTICS - ANOVA (Experimental) - Two way additive, Independent (Y ~ A + B) | FDR threshold to include in results | N/A | N/A | N/A | N/A | N/A | N/A |
| STATISTICS - ANOVA (Experimental) - Two way additive, Independent (Y ~ A + B) | Choose variable to test | N/A | N/A | N/A | N/A | N/A | N/A |
| STATISTICS - ANOVA (Experimental) - Two way additive, Independent (Y ~ A + B) | Choose grouping variables | N/A | N/A | N/A | N/A | N/A | N/A |
| STATISTICS - ANOVA (Experimental) - Two way additive, Independent (Y ~ A + B) | Choose variables for modeling | N/A | N/A | N/A | N/A | N/A | N/A |
| STATISTICS - ANOVA (Experimental) - Two way additive, Independent (Y ~ A + B) | Ind V1 | N/A | N/A | N/A | N/A | N/A | N/A |
| STATISTICS - ANOVA (Experimental) - Two way additive, Independent (Y ~ A + B) | Ind V2 | N/A | N/A | N/A | N/A | N/A | N/A |
| STATISTICS - ANOVA (Experimental) - Two way additive, Independent (Y ~ A + B) | Choose members to include | N/A | N/A | N/A | N/A | N/A | N/A |
| STATISTICS - ANOVA (Experimental) - Two way Interaction (Y ~ A * B) | FDR threshold to include in results | N/A | N/A | N/A | N/A | N/A | N/A |
| STATISTICS - ANOVA (Experimental) - Two way Interaction (Y ~ A * B) | Choose variable to test | N/A | N/A | N/A | N/A | N/A | N/A |
| STATISTICS - ANOVA (Experimental) - Two way Interaction (Y ~ A * B) | Choose grouping variables | N/A | N/A | N/A | N/A | N/A | N/A |
| STATISTICS - ANOVA (Experimental) - Two way Interaction (Y ~ A * B) | Choose variables for modeling | N/A | N/A | N/A | N/A | N/A | N/A |
| STATISTICS - ANOVA (Experimental) - Two way Interaction (Y ~ A * B) | Ind V1 | N/A | N/A | N/A | N/A | N/A | N/A |
| STATISTICS - ANOVA (Experimental) - Two way Interaction (Y ~ A * B) | Ind V2 | N/A | N/A | N/A | N/A | N/A | N/A |
| STATISTICS - ANOVA (Experimental) - Two way Interaction (Y ~ A * B) | Choose members to include | N/A | N/A | N/A | N/A | N/A | N/A |
| STATISTICS - ANOVA (Experimental) - Repeated measures, Between subjects interaction ($Y \sim A * B + \text{Error}(C)$) | FDR threshold to include in results | N/A | N/A | N/A | N/A | N/A | N/A |
| STATISTICS - ANOVA (Experimental) - Repeated measures, Between subjects interaction ($Y \sim A * B + \text{Error}(C)$) | Choose variable to test | N/A | N/A | N/A | N/A | N/A | N/A |
| STATISTICS - ANOVA (Experimental) - Repeated measures, Between subjects interaction ($Y \sim A * B + \text{Error}(C)$) | Choose grouping variables | N/A | N/A | N/A | N/A | N/A | N/A |
| STATISTICS - ANOVA (Experimental) - Repeated measures, Between subjects interaction ($Y \sim A * B + \text{Error}(C)$) | Choose variables for modeling | N/A | N/A | N/A | N/A | N/A | N/A |
| STATISTICS - ANOVA (Experimental) - Repeated measures, Between subjects interaction ($Y \sim A * B + \text{Error}(C)$) | Variable 1 | N/A | N/A | N/A | N/A | N/A | N/A |
| STATISTICS - ANOVA (Experimental) - Repeated measures, Between subjects interaction ($Y \sim A * B + \text{Error}(C)$) | Variable 2 | N/A | N/A | N/A | N/A | N/A | N/A |
| STATISTICS - ANOVA (Experimental) - Repeated measures, Between subjects interaction ($Y \sim A * B + \text{Error}(C)$) | Subject | N/A | N/A | N/A | N/A | N/A | N/A |
| STATISTICS - ANOVA (Experimental) - Repeated measures, Between subjects interaction ($Y \sim A * B + \text{Error}(C)$) | Choose members to include | N/A | N/A | N/A | N/A | N/A | N/A |
| HEATMAP AND OUTLIER DETECTION - Heatmap | Heatmap parameters | N/A | N/A | N/A | N/A | N/A | N/A |
| HEATMAP AND OUTLIER DETECTION - Heatmap | Choose labels | N/A | N/A | N/A | N/A | N/A | N/A |
| HEATMAP AND OUTLIER DETECTION - Heatmap | Kmeans k-value | N/A | N/A | N/A | N/A | N/A | N/A |
| HEATMAP AND OUTLIER DETECTION - Heatmap | cutree row clusters | N/A | N/A | N/A | N/A | N/A | N/A |
| HEATMAP AND OUTLIER DETECTION - Heatmap | Cutree col clusters | N/A | N/A | N/A | N/A | N/A | N/A |
| HEATMAP AND OUTLIER DETECTION - Heatmap | Clustering method (R hclust) | N/A | N/A | N/A | N/A | N/A | N/A |
| HEATMAP AND OUTLIER DETECTION - Heatmap | Column distance method | N/A | N/A | N/A | N/A | N/A | N/A |
| HEATMAP AND OUTLIER DETECTION - Heatmap | Row distance method | N/A | N/A | N/A | N/A | N/A | N/A |
| HEATMAP AND OUTLIER DETECTION - Heatmap | Heatmap colors | N/A | N/A | N/A | N/A | N/A | N/A |
| HEATMAP AND OUTLIER DETECTION - Heatmap | Choose filtering column for heatmap | N/A | N/A | N/A | N/A | N/A | N/A |
| HEATMAP AND OUTLIER DETECTION - Heatmap | Filtering stat direction | N/A | N/A | N/A | N/A | N/A | N/A |
| HEATMAP AND OUTLIER DETECTION - Outlier detection | Alpha value | N/A | N/A | N/A | N/A | N/A | N/A |
| HEATMAP AND OUTLIER DETECTION - Outlier detection | k value | N/A | N/A | N/A | N/A | N/A | N/A |
| HEATMAP AND OUTLIER DETECTION - Outlier detection | KNN searchtype | N/A | N/A | N/A | N/A | N/A | N/A |
| HEATMAP AND OUTLIER DETECTION - Outlier detection | Proportion of possible outliers | N/A | N/A | N/A | N/A | N/A | N/A |
| HEATMAP AND OUTLIER DETECTION - Outlier detection | Sample size to calulate empirical threshold | N/A | N/A | N/A | N/A | N/A | N/A |
| COLOCALIZATION ANALYSIS | Number of colocalized features? | N/A | N/A | N/A | N/A | N/A | N/A |
| COLOCALIZATION ANALYSIS | Sorted by | N/A | N/A | N/A | N/A | N/A | N/A |

